# A behavioral roadmap for the development of agency in the rodent

**DOI:** 10.1101/2023.11.10.566632

**Authors:** C Mitelut, M Diez Castro, RE Peterson, M Gonçalves, J Li, MM Gamer, SRO Nilsson, TD Pereira, DH Sanes

**Author notes:** **Materials & Correspondence:** Mitelut C. Equal contribution first author (C Mitelut, M Diez Castro). Co-last authors (TD Pereira, DH Sanes).

## Abstract

Behavioral interactions within the nuclear family play a pivotal role in the emergence of agency: the capacity to regulate physiological, psychological and social needs. While behaviors may develop over days or weeks in line with nervous system maturation, individual behaviors can occur on sub-second time scales making it challenging to track development in lab studies with brief observation periods, or in field studies with limited temporal precision and animal identification. Here we study development in families of gerbils, a highly social rodent, collecting tens of millions of behavior time points and implementing machine learning methods to track individual subjects. We provided maturing gerbils with a large, undisturbed environment between postnatal day 15 and the age at which they would typically disperse from the family unit (day 30). We identified complex and distinct developmental trajectories for food and water acquisition, solitary exploration, and social behaviors, some of which displayed sex differences and diurnal patterns. Our work supports the emergence of well-delineated autonomous and social behavior phenotypes that correlate with specific periods and loci of neural maturation.

## INTRODUCTION

At the center of mammalian behavior development lies the emergence of agency (Sokol et al 2015): the capacity to self-organize one’s actions towards numerous survival and thriving goals (Mahner and Bunge, 1997, Sokol et al 2015). Agency development thus involves the onset of food and water foraging, learning through environmental affordances (Gibson 1982), regulation of emotions for selecting complex goals and acting intentionally (Lerner et al 2015), and-in humans-sharing and creating actions based on cultural and social norms (Lewis 2014; Mayes & Lewis 2012).

Studying behavior across species has long been appreciated (Beach, 1950) especially where human correlates of neurodevelopmental sequences are known (Worman et al 2013). In rodents, for example, the development of agency commences early within the nest with thermoregulation (Shelton and Alberts 2018) or food odor orientation (Al Ain et al 2011), increases with exploration outside the nest, and culminates with dispersal during which family social bonds are severed (Howard 1949; Howard 1960; Gerlach 1990; Gerlach 1996; Groó et al 2013; Wang et al 2017). These behavioral changes overlap substantially with the maturation of sensory (eye/ear development: Altman & Sudarshan 1975; sniffing: Westerga & Gramsbergen 1990), motor (Blumberg & Adolph 2023) and spatio-cognitive systems (Wills et al 2013). Social and environmental factors also drive development, including an attraction of pups to adults that are outside of the nest (Galef & Clark 1971; 1972; Alberts & Leimbach 1980), a decline in nest signals that promote affiliation (Grota & Ader 1969; 1970; Levin & Stern 1975; Goodwin & Yacko 2004), and instrumental learning about social variables (Chen & Amsel 1980). The development of agency in rodents thus requires understanding how complex behaviors emerge from an interplay of sensory-motor-cognitive system maturation and interactions with parents and siblings.

Social repertoires differ dramatically among social species, with some exhibiting much greater social structures. Mongolian gerbils, in particular, are a favorable species for the study of family development as they exhibit favorable behaviors such as monogamous pair-bonding (Tchabovsky et al 2019), family member division into founder pairs and integrating or expelling of family members (Scheibler et al 2004), dynamic association strategies to reflect life histories (e.g., during food-hoarding season; Deng et al 2017), coordination between male and female parents during pup rearing (Ostermeyer & Elwood 1984), female mating preferences for relatedness (Valsecchi 2002), increased prosociality towards pairbond partners-but not others-following testosterone boosts (Kelly et al 2022), and gradual expulsion of males with increased reproductive capacities from family group (Scheibler et al 2006). In the wild, gerbil pups largely remain in the nest until postnatal day (P) 16, and while they increasingly leave the nest from P18-P22 (Kaplan & Hylan 1972), they remain attracted to maternal nest odors until at least P42 (Yahr & Anderson-Mitchell 1983). During this time they do not increase vocalizations when the mother is absent from the nest, and receive a similar amount of care from the father (Ostermeyer & Elwood 1984). This period of significant development-from complete dependency on the mother until ∼P15 to complete independence by P30-P40-should contain a large array of complex behavior interactions, yet the interaction of individual behaviors is uncertain. Critically, characterizing all the behavioral stages and neural mechanisms that yield independent animals with full agency-requires an approach that combines the advantages of field studies with the precise identity and subsecond precision tracking available during transient lab observations (Lin & Wilbrecht 2022; Bordes et al 2023).

To gain new insights into how autonomous and social behaviors develop within family groups we used uninterrupted home-cage video recordings in concert with a machine learning framework to continuously track individual Mongolian gerbil family members (2 adults and 4 pups) for two weeks. We began recording when animals first leave the nest (∼P15) and collected > 1000 hours of video data (∼100 million video frames) across three cohorts of animals, and implemented and adapted pose estimation tools (Pereira et al 2022), behavior classifiers (Nilsson et al 2020) and custom algorithms for behavior classification. Our results reveal distinct timelines for the emergence of social and nonsocial behaviors, sex-based differences in the behaviors of both adult and developing pups, and the emergence of diurnal behaviors. We propose that over these pivotal two weeks of development social and nonsocial measures of agency display distinct time courses, likely supported by complex interactions of brain maturation and novel social and environmental affordances. Our work contributes a novel framework with which to appraise quantitative studies of central nervous system (CNS) maturation.

## RESULTS

### Tracking behaviors of individual gerbil family members in home-cage environments

*Capturing individual gerbil behavior across weeks with subsecond temporal precision.* We obtained video recordings from three gerbil family cohorts from postnatal days (P) 15-30 (Fig 1a; see also Methods). Prior to data collection, all gerbils were shaved with a distinct pattern to facilitate identification of individuals by animal tracking machine vision algorithms (Fig 1b). Families were placed in an isolated enriched home environment at P14, and we began data collection on P15, the age at which gerbil pups begin to leave their nest (Ågren 1989). Data collection ended on P30, the age at which juveniles are weaned in our colony (Fig 1c). We captured more than 30 million frames of video data per cohort across three family cohorts with varying numbers of female and male pups (Fig 1d).

**Fig 1.**
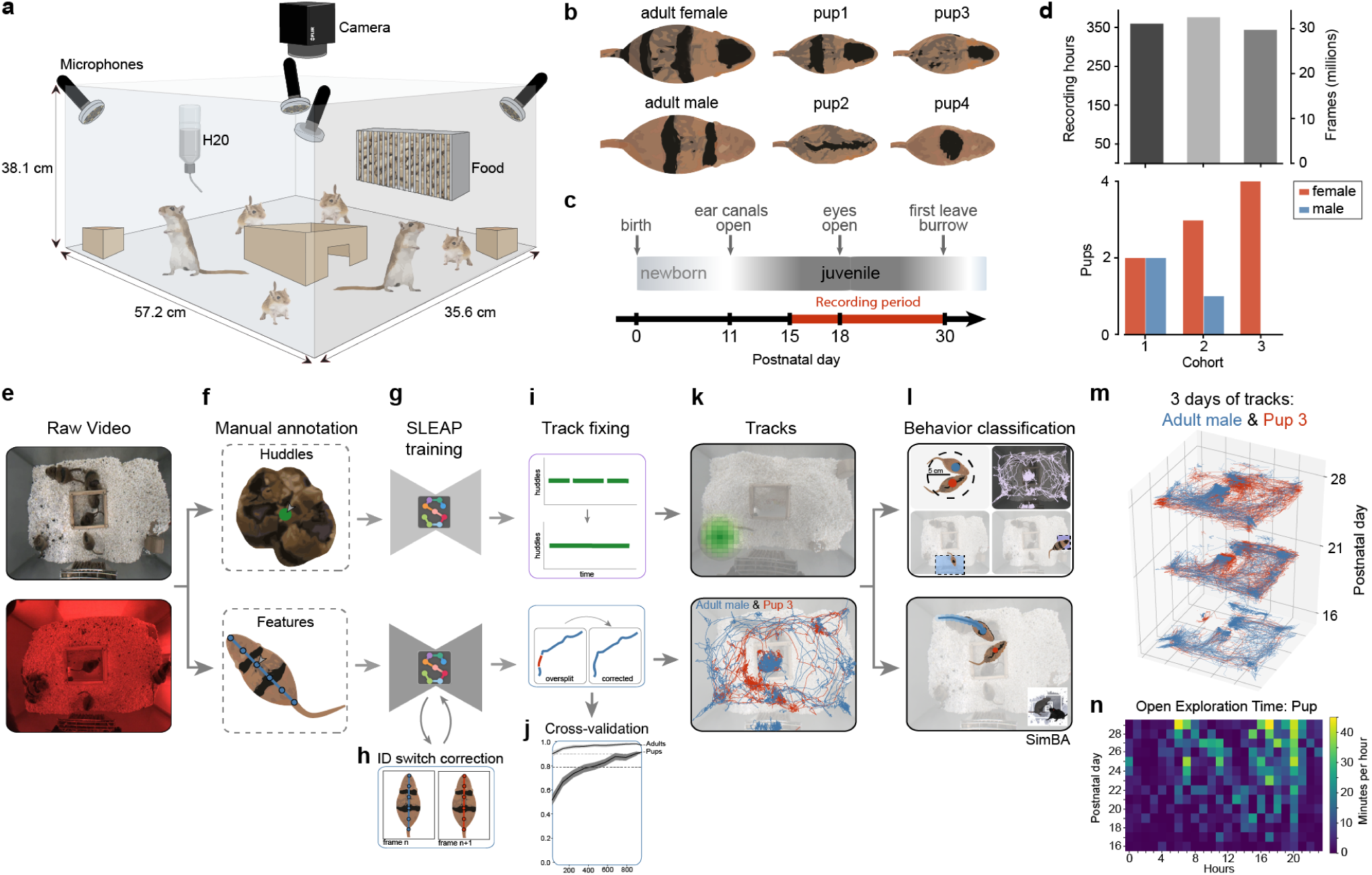
Capturing individual gerbil behavior across weeks of development with subsecond precision. **(a)** Homecage and recording setup using continuous, synchronized video (single overhead camera at 24 fps) and audio (4 ultrasonic microphones at 125 kHz) recordings in environments with *ad libitum* access to food and water and enrichment consisting of one wooden hut and two blocks. (**b)** Individual gerbil identification based on distinct shaving patterns. **(c)** Recording periods of 15 days from postnatal (P) day 15 to 30. (**d)** Top: number of recording hours and frames per cohort; Bottom: number of female and male pups per cohort (bottom). (**e-l):** Workflow of animal tracking and behavior analysis. (**e)** Acquisition of raw video in a 12 hr light / dark (infrared) cycle. (**f)** Manual annotation of two categories: huddles (2 or more gerbils huddling together in the nest area) and features (individual animals with a 6-node skeleton). (**g)** Each light condition (light and dark) and cohort required separate training resulting in six feature models and six huddle models. (**h)** Method for identification of the identity switches between consecutive frames. (**i)** Method for interpolating small missing tracks based on id-swapping cues. (**j)** Cross-validation of model performance (note: chance is 1/6 = 0.17). (**k)** Visual representation of nest location and animal tracks. (**l)** Top: sketch of behaviors classified using heuristics (e.g. time spent in a region of interest and distance traveled); Bottom: sketch of approach behaviors captured using SimBA classifier. (**m)** Two animal tracks (adult male and pup) across 3 days of development. (**n)** Example ethogram of open exploration time per hour for one pup across 15 days of development.

*Tracking individual gerbils behaviors.* We used SLEAP (Pereira et al 2022) to identify individual gerbils and gerbil huddles across day and night cycles (Fig 1e; see also Methods). For tracking huddles, we trained SLEAP on manually annotated frames with groups of 2 or more gerbils that were huddling at a stable location, and for tracking gerbils in the open field we annotated 6 body features along the spine of the gerbils on randomly chosen 1000 frames from dozens of videos (Fig 1f). Because of idiosyncratic differences in shaving patterns and light conditions across the cohorts, we trained separate models for each cohort on day and night conditions, resulting in 6 body-feature models and 6 huddle models (Fig 1g).

We additionally developed methods for improving animal identity tracking: (i) an identification-switch detection pipeline that automatically located identity tracking errors on sequential video frames (Fig 1h; see Methods), (ii) a track fixing pipeline that interpolated short missing periods of huddling or individual tracks (Fig 1i, top) and (iii) an identity swap pipeline that detected and corrected obvious track identity swaps (Fig 1i, bottom). We used 10-fold cross validation to evaluate SLEAP *identity* accuracy and found a true positive rate of 0.97 (0.04 std) for adult animals and 0.91 for pups (0.05) (Fig 1j). Errors in *feature* (i.e. swapping of body parts within an animal) were extremely low (< 0.1 %) and we additionally used centroids to represent animal locations which are less sensitive to such errors (see Methods). Using our pipelines, we were able to track the huddling (Fig 1k, top) and open field behaviors of multiple animals across weeks of continuous recording (Fig 1k, bottom).

*Classifying animal behavior.* We implemented two types of methods for classifying behavior: heuristic-based methods and supervised classification methods (Fig 1l). For heuristic-based classifications we used Regions-of-Interest (ROIs) to identify when animals were near the food hopper or water spout regions, and proximity metrics for detecting when gerbils were within 5 cm of one another (Fig 1l, top; see Methods). For approach behaviors we used a customized version of SimBA (Nilsson et al 2020) to train a classifier to capture approach behaviors from manually labeled examples (Fig 1l, bottom) (see Methods). Using our pipelines, we were able to track individual gerbils over periods of two weeks with subsecond predictions (Fig 1m) and generate ethograms that captured postnatal day and hourly dynamics over several behaviors (Fig 1n for an example).

In sum, we found that unique shaving patterns, and tracking algorithms with customized algorithms enabled the tracking of multiple individuals over weeks of behavior. This enabled us to characterize the development of individual and group behaviors in gerbil pups during the period of development when animals adopt independent repertoires (See results below).

### The developmental trajectories of biophysical agency

Using the processing pipelines and heuristics described above, we evaluated behavior development in gerbil pups over two week periods (P16 to P29 inclusive) across three cohorts of animals (Fig 2). We first focused on the development of biophysical agency, i.e. the development of capacities to regular and sustain oneself *independently* to other animals’ behaviors including the adult parents.

**Fig 2.**
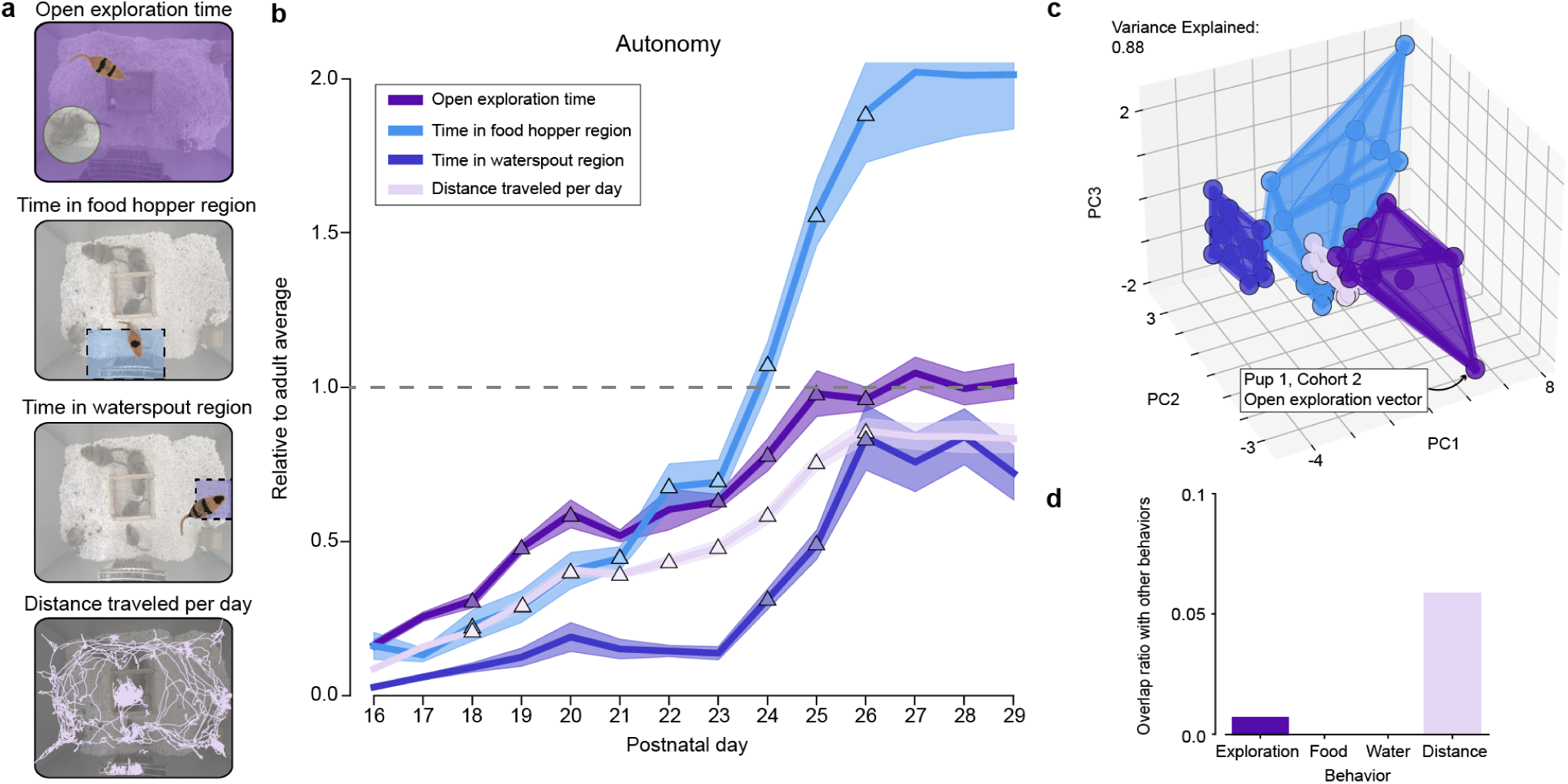
The development of autonomous behaviors. **(a)** Sketches illustrating autonomous behaviors including open exploration time, exploration of food hopper and waterspout regions, and total distance traveled. (**b)** Average environment exploration trajectories in gerbil pups relative to the adult average on P29 (lines) and standard error (SEM; shading) with indication of rapid development days (triangles). **(c)** Convex hull overlap analysis of individual animal multi-day trajectories with intersections computed in 3D PCA space (i.e. PCs 1-3) with an example of a single pup trajectory for open exploration time. (**d)** Overlaps in PCA space from (c) show limited inter-behavior overlap.

*Diverse and dynamic development of territory exploration in gerbil pups.* We tracked the amount of time gerbils spent exploring their habitat outside the nest, in the food hopper region, and in the water spout region, as well as the total distance traveled per day (Fig 2a; see also Methods). As we were interested in the maturation of behavior, we evaluated pup behavior relative to the average adult behavior recorded when pups were P29 (Fig 2b). We selected P29 as the adult reference date as we found behaviors of adults stabilized by this period. We differentiated between periods during which behaviors were (i) *stable* (Fig 2b-colored lines) or (ii) *rapidly developing* (Fig 2b-triangles). For each behavior, we defined a postnatal day as a period of *rapid development* when it was statistically different from the previous 2 days and the following day (2-sample K-S test; p-value < 0.05; see also Methods).

We found that pup *distance traveled per day* started at 8.72 ± 1.71 % of adult levels (or 8.01 ± 1.53 meters/day) and increased continuously until P26 where it stabilized near adult levels at 85.90 ± 13.08 % (or 78.98 ± 9.91 meters/day). Pup *open exploration time* followed a similar trajectory, starting at 16.34 ± 5.48 % (or 0.88 ± 0.32 hrs/day) and reaching 96.16 ± 12.82 % (or 5.17 ± 0.84 hrs/day) (Fig 2b). We note that both distance traveled and environment exploration time underwent rapid development for most of the days between P17 and P25. These open environment exploration findings suggest a steady and continuous maturation rate with pup behavior becoming similar to adults by P26, 11 days after gerbil pups leave the nest.

In contrast, food hopper exploration increased slowly from P16 at 16.12 ± 14.59 % of adult levels (or 0.14 ± 0.10 hrs/day) to P21 at 44.54 ± 24.92 % (or 0.44 ± 0.13 hrs/day). They underwent a further period of rapid development between P22 until P26 stabilizing from P27-P29 at 201.58 ± 68.18 % (or 1.92 ± 0.25 hrs/day), more than 100% higher than adult averages (Fig 2b). This suggests that in addition to learning the nutritional value of the food hopper region, pups may also value this space as a socialization space significantly more than adults (see more below).

Water spout region exploration also displayed three stages, but with a much later trajectory than food: (i) an initial stage with pups spending less than 10% of the adult averages from P16 at 2.75 ± 1.44 % (or 0.007 ± 0.004 hrs/day) to P23 at 13.75 ± 7.16 % (or 0.04 ± 0.02 hrs/day), (ii) a period of rapidly increasing water spout access from P24 at 31.28 ± 11.10 % (or 0.07 ± 0.02 hrs/day) to P26 at 83.68 ± 35.12 % (or 0.19 ± 0.07 hours), and (iii) a period of stability from P27 to P29 where gerbil pups spend 77.27 ± 30.26 % relative to the adults (or 0.18 ± 0.02 hrs/day). The rapid increase from P24 to P26 may reflect a discovery of water sources and a rapid reduction in the time spent nursing.

*Distinct inter-behavior developmental trajectories of territory exploration in gerbil pups.* We next investigated whether individual pup behaviors were distinct from one another. We developed a method that computed the spatial overlap of each of the behaviors by using principal component analysis (PCA) first to reduce the dimensionality of the multi-day behavior trajectories and then computed the convex hull overlaps across behaviors (Fig 2c; see Methods). We used 3 PCs to compute the (3d) intersection of the behaviors as this provided approximately 80% (or more) variance explained for all the behaviors (see Methods). We found this method was more interpretable than Kullback-Leibler divergence or Mahalanobis distances which do not directly quantify overlap as a percentage. Across the four behaviors tracked we found only minimal overlap between open exploration time (0.73%) and distance traveled per day behaviors (5.9%). This confirmed our qualitative observations that open exploration time and distance traveled per day trajectories were most similar to each other but that overall, environmental exploration behaviors had distinct trajectories.

In sum, we found that across four behavioral measures we could identify distinct developmental time courses that were significantly different from each other (and adults). Thus, for example, food and water consumption develop on different timelines and have their own unique periods of rapid change-suggesting complex interactions between drives for nutritional needs and independent exploration of the environment.

### The developmental trajectories of animal pair social states

We next sought to characterize pair-wise social behaviors to capture dependencies between individual family members and evolving pup-pup relationships (Fig 3). We defined social behaviors as those that *involved another animal* such as time spent huddling in the nest or approaches between two animals.

**Fig 3.**
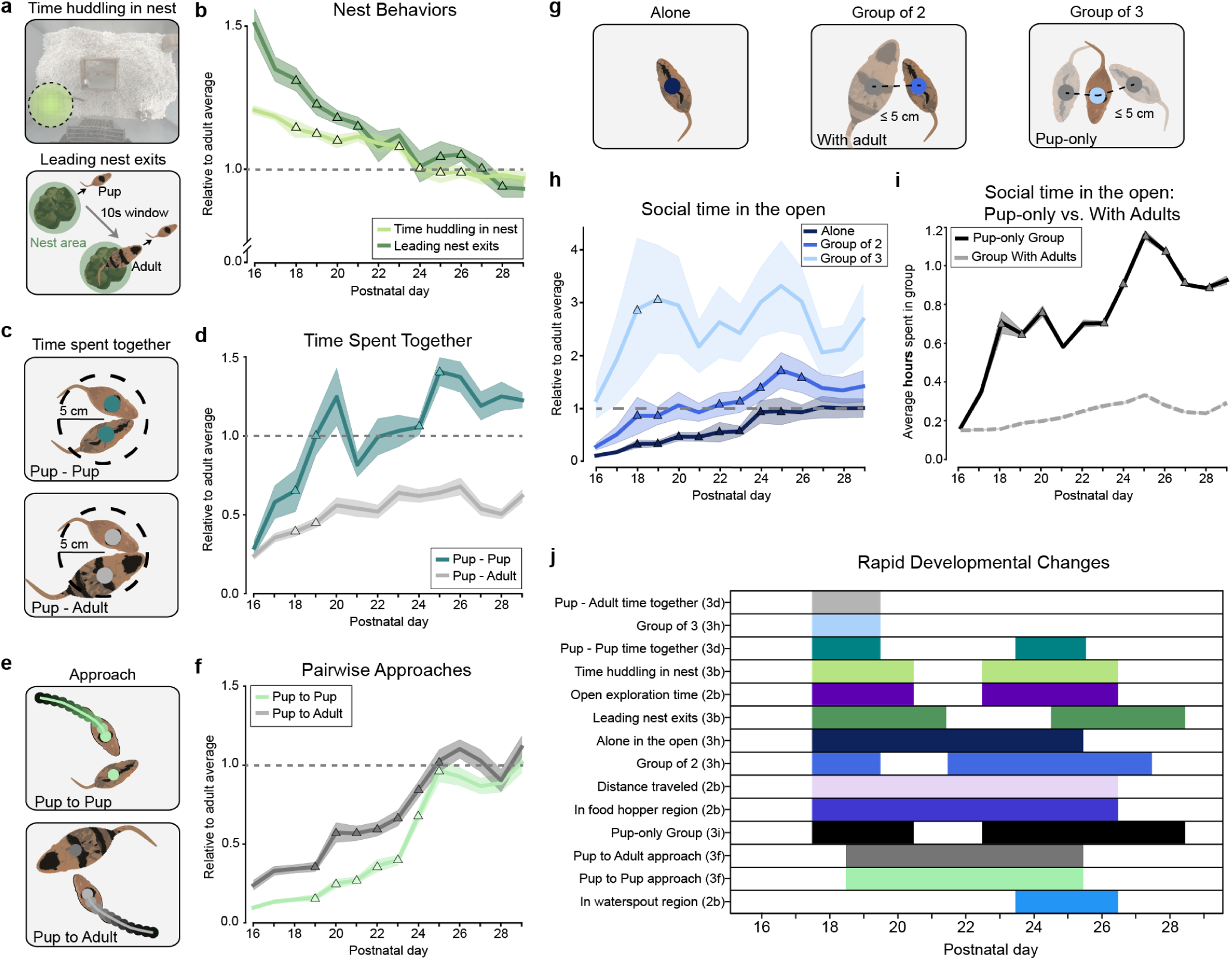
The development of psycho-social driven behaviors. **(a)**. Sketches illustrating nest behaviors (time huddling in nest) and sequences of adults following pups out of the nest (within 10sec). **(b).** Average huddling time in the nest (light green) and leading nest exits (dark green) relative to the adult average on P29 (shading-SEM). **(c)** Sketch illustrating pairwise proximity behaviors. **(d)** The average time of pup time spent in close proximity with each other (turquoise) and with the adults (gray) relative to the adult average on P29 (shading-SEM). **(e)** Sketches of pairwise approaches. **(f)** The average number pup approaches towards other pups (green) and adults (dark gray) relative to the adult average on P29 (shading-SEM). **(g)** Sketches of single or multi group proximity. **(h)** Average time spent in the open by pups as individuals (dark blue) or groups of two (blue) or groups of three (light blue) relative to the adult average on P29 (shading-SEM). **(i)** Average hours spent in groups with pups (black) vs. groups with adults (dashed gray line). (**j)** Summary of rapid development periods (colors) across all behaviors tracked.

*The development of pairwise social interactions in gerbil pups.* With respect to *nest and huddling behaviors*, we computed two metrics: (i) time that pups spent huddling in the nest; and (ii) amount of times adults followed pups out of the nest-to capture the tendency of adults to retrieve or care for younger pups (Fig 3a; see also Methods). We found that pups spent more time than adults in the nest on P16 at 121 ± 2 % (or 23.12 ± 0.02 hrs/day) and this value reached and stabilized at the adult average by P24 at 104 ± 6 % (or 19.80 ± 1.05 hrs/day). Similarly, pups were followed out of the nest by parents significantly more than adults on P16 at 151 ± 0.17 % (and overall followed 76.09 ± 8.57 % of the time), and this nest following behavior stabilized also around P24 at 107 ± 12 % (or 53.53 ± 5.88 %) (Fig 3b) (Note: the two behaviors had similar developmental trajectories with nest exits having higher variance and only 0.06 overlap where as and nest huddling having very low variance and 0.46 overlap).

With respect to *pairwise social exploration,* we tracked the amount of time pups spent together with each other or with other adults (Fig 3c). We found that pups tended to spend significantly more time near pups than adults (Fig 3d). In particular, at P16 pup-pup pairwise exploration was 29.23 ± 4.41 % (or 9.15 ± 4.15 minutes/day), but increased rapidly to adult levels within a few days reaching 100.48 ± 11.89 % (or 28.03 ± 7.55 minutes/day) by P19 and stabilizing on P29 at 122.80 ± 4.74 % (or 33.67 ± 1.51 minutes/day). In contrast, adult-pup pairwise exploration time started on P16 at 24.31 ± 2.35 % of the adult average (or 5.35 ± 1.88 minutes/day) and increased slower reaching a peak around P26 at 68.13 ± 5.30 % (or 16.02 ± 4.03 minutes/day). These findings suggest that pups interacted significantly more with other pups throughout development as compared to adult interactions (e.g. pup-pup interactions were on average 193.2% of pup-adult interactions overall days). Note there was no overlap in the PCA space (not shown).

For *approach* behaviors, we found a similar pattern between pup-pup and pup-adult approaches with some important differences (Fig 3e). Pup-pup approaches were ∼3-fold lower on P16 at 9.52 ± 7.96 % (or 3.52 ± 2.97 approaches/day), as compared to pup-adult approaches at 23.77 ± 15.28 % (or 10.08 ± 6.38 approaches/day) (Fig 3f). Both types of approaches increased gradually, with pup-pup approaches undergoing rapid development from P20 at 24.70 ± 13.35 % (or 11.14 ± 5.44 approaches/day) to P25 at 96.23 ± 23.47 % (or 39.76 ± 16.21 approaches/day) before leveling off to near adult levels: P26 to P29 averages 92.24 ± 17.54 % (or 38.29 ± 13.29 approaches/day). Pup-adult approaches also underwent rapid development from P20 at 57.07 ± 22.18 % (or 28.21 ± 8.95 approaches/day) to P25 at 102.50 ± 21.26 % (or 35.67 ± 9.40 approaches/day) and leveling off to adult levels (P26-P29: 103.51 ± 32.10 %, or 44.18 ± 12.73 approaches/day for pup-adults, compared to 43.33 ± 9.50 for adult-adults). The convex hull overlap between pup-pup and adult-pup approaches was ∼10% (not shown) suggesting only limited similarity between how individual pups approached adults vs. other pups. Taken together with the pairwise socialization times, the pairwise approach results suggest while pups approach other pups fewer times per day during early development (Fig 3f) they spend significantly more time together following such approaches (Fig 3d) possibly indicating different reasons or motivations for the two types of interactions.

Lastly, we computed *group socialization* time as the time pups spent in groups of two or three gerbils (Fig 3g). We found that pups socialized in groups of three gerbils up to 3 fold more than adults did and that this behavior was present as early as P18, approximately 2 days after pups left the nest (Fig 3h). Looking specifically at the absolute time spent in groups with pups vs. adults, we found that beginning with P18 pups spent a factor of 2-3 more time in groups with pups than in groups with adults (Fig 3i). This establishes again a significant drive for socialization for pups with other pups-rather than being monolithic across all ages. Note there was no significant overlap of convex hulls in PCA space (not shown).

*Complex and heterogeneous development periods.* Over all behaviors studied, we found distinct sequences of rapid development and stabilization periods. When taken together (Fig 3j), biophysical agency (e.g., food and water acquisition) appeared to have the latest onset, occurring after P21. In contrast, many social behaviors (groups of 2 or 3) tended to have an early onset with a later period of maturation. The delay of nutrition-seeking behaviors are perhaps explained by Mongolian gerbils having a longer period of parental care and integrity as a multigenerational family We also note that rapid development after P26 stopped for all but large group socialization suggesting that pup development across several axes we studied was largely complete, at least given the limitations of a closed environment.

In sum, we found that most environmental exploration and social behaviors had unique developmental trajectories with intra-behavior differences being substantially greater than inter-animal differences. This supports the unfolding of different classes of behavior on distinct time courses correlating with the maturation of several components of the nervous system in rodents between P15 and P30 (see Discussion). We also found that there are distinct rapid development periods during which pup behavior changed rapidly over a period of a few days or longer. This suggests that in addition to distinct behavioral correlates to the maturing central nervous system, there are narrow periods of a few days during which specific behaviors mature significantly.

### The development of multi-animal social state dynamics

We next evaluated the development of multi-animal social behaviors by focusing on the complexity of social states and configurations in the home cage. We were particularly interested in quantifying the complexity of multi-animal social configurations in the space of their environment. We partitioned the cage floor into a 5 by 4 grid and defined a social state at each recording time point as the number of animals at each location (Fig 4a,b) (see also Methods). This enabled us to generate a network-based analysis where each time point in our recording generated a 20 Dimensional (20D) social state vector and we could generate a network connectivity matrix by connecting all social states that occurred sequentially (Fig 4c; see also Methods). The network connectivity matrix enabled us to generate graphs that represented the social state dynamics in our cohorts (Fig 4d).

**Fig 4.**
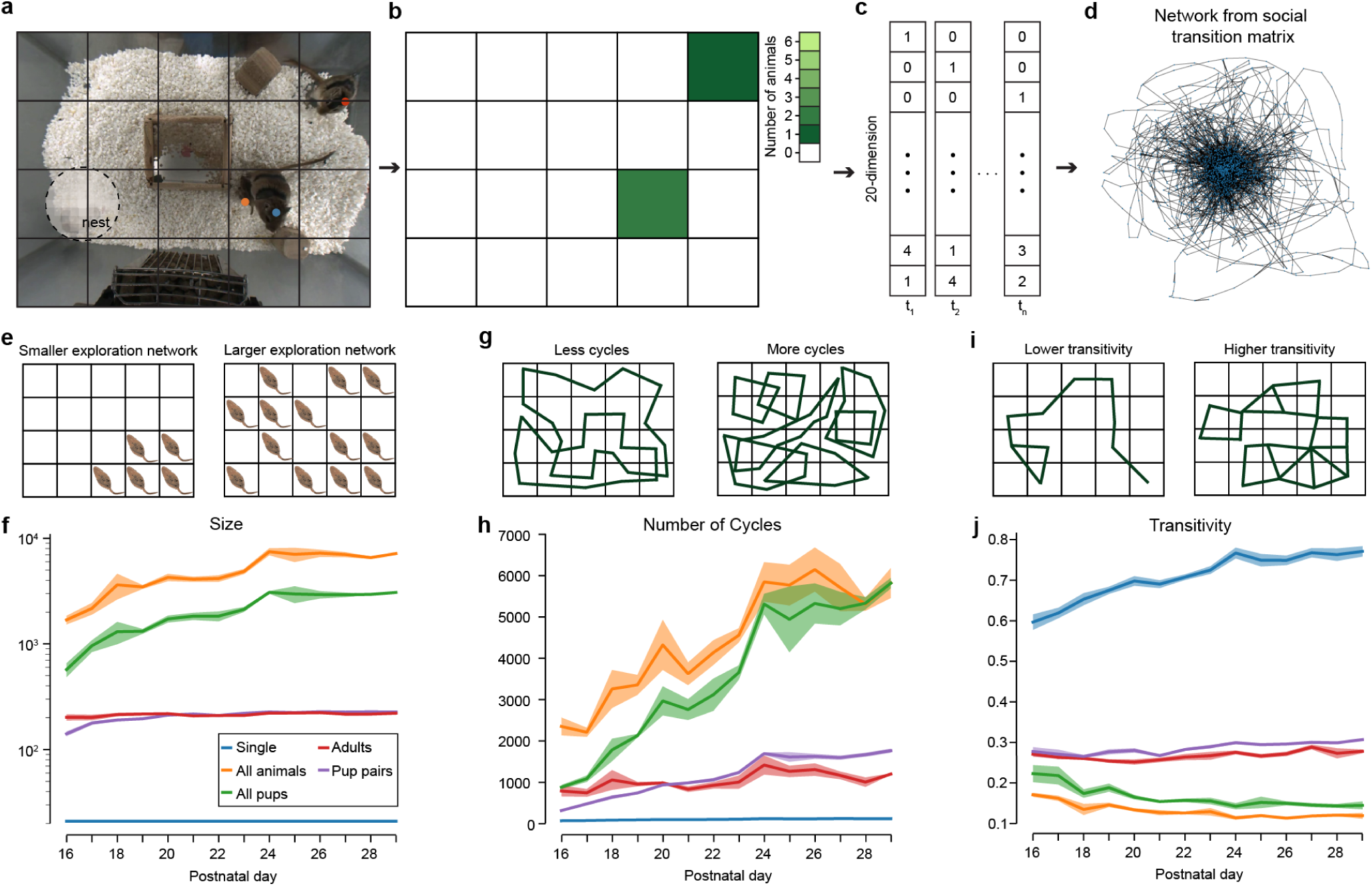
The development of social behavior complexity. **(a)** Partitioning the home cage environment into a 5 x 4 social-state grid. **(b)** Binarization of social states in (a) by the number of gerbils in each grid square. **(c)** Converting social-states to sequences of 20D vectors (note: vectors do not correspond to panel (b)). **(d)** Network graph obtained from social-state transitions for 1 day. **(e)** Sketch depicting behavior of a single gerbil that would generate smaller vs. larger sized networks (note: multi-animal configurations have more than 2 dimensions and are not simple to visualize; see Methods). **(f)** Network sizes vs development for single and different groups of gerbils (note logarithmic scaling of y-axis). **(g)** Sketch depicting behavioral paths of a single gerbil that would generate graphs with less vs. more cycles. **(h)** Number of graph cycles vs development. **(i)** Sketch depicting path of a single gerbil that would generate graphs with lower vs. higher transitivity. **(j)** Transitivity metrics vs. development.

*The number of social-state configurations increases across development for groups of animals.* We computed a social-state graph for single animals, pairs (2 animals), all pups (4 animals) and all animals (6 animals) for each PDay of our study (Fig 4e,f). We first evaluated graph sizes which represent the number of unique social configurations in each graph (4e). We found that between P16 and P19, the graph sizes that grew most significantly were all-pup graphs: P29/P16 ratio = 5.37, and all-animal graphs: P19/P16 ratio = 4.24 (Fig 4f). In contrast, social state complexity for pup-pup graphs increased much less: P29/P16 ratio = 1.61, and adult-adult complexity was mostly stable over time: P29/P16 ratio = 1.09. This suggests that social behavior complexity increased the most for larger groups of animals in ways possibly not accounted for solely by pair-wise social complexity increases alone. Additionally, consistent with our findings for social states in animal-pairs, we found that social state networks for larger animal groups (i.e. 4 and 6) leveled off around P24 (Fig 4f).

*The number of social-state paths increases across development for groups of animals but not adult pairs.* We next computed the number of cycles for each day and animal grouping (Fig 4g,h). The number of cycles can be viewed as the number of unique paths in the social state space that animals can traverse over time (Fig 4g). We found that both all-pup and all-animal groupings showed a substantial increase in the number of cycles over development up to P24 when we observed qualitative leveling off as for network sizes (all-pup: P29/P16 ratio = 6.54; all-animal: P29/P16 ratio = 2.47). Similarly, pup-pup pairwise social states had a significant increase over development: P29/P16 ratio = 5.44. In contrast, the adult-adult social states had very similar numbers of cycles across development: P29/P16 ratio = 1.51. Interestingly, we found the average path length for all groups to be relatively stable over development or slightly decreasing (not shown) suggesting that individual social-state dynamical paths were not getting significantly larger-but rather were increasing in number, likely driven by new areas and configuration to explore. Taken together, the developmental dynamics of the number of cycles suggest that novel social-state configurations are generated during the period of P16 to P24 and are driven by pup development with less contribution from adults.

*Social exploration increases across development for large groups but not pairs or single animals.* We lastly computed the transitivity of each social configuration network (Fig 4i,j). Transitivity refers to the overall probability of a node to have interconnected adjacent edges (i.e. the ratio of possible triangles in the graph) (Fig 4i). In our paradigm, transitivity can be viewed as how comprehensively each social configuration is explored by the animals. High transitivity values would indicate that gerbils spend more time in individual configurations with slight alteration-whereas low transitivity values indicate that gerbils transition through more diverse social configurations without spending time exploring individual configuration. Put in simpler terms, high transitivity could be viewed as a *tendency to exploit* locally specific social configurations whereas low transitivity could be viewed as a *tendency towards exploring* more novel and different social configurations.

We found that single animals showed an increase in transitivity over time: P29/P16 ratio = 1.29-suggesting a tendency for single animals to exploit existing paths rather than pursuing new ones. In contrast, pairs of animals (adults and pups) had slight tendencies for exploration: pup pairs P29/P16 = 1.10; and adult pairs P29/P16 = 1.03. In contrast, all-pup and all-animal groups had significant trends towards increased exploration of novel social configurations: all pups P29/P16 = 0.65; and all animals P29 / P16 = 0.70. Taken together, these results suggest that when in larger groups animals tended to seek increasingly novel types of social-state configurations, essentially exploring the social state space more. Whereas as individuals or pairs, they did so much less. This suggests that increased group sizes could be supportive of environmental exploration and that single animals or smaller groups tend to use more stereotyped or repeated trajectory patterns.

### Sex-based behavioral difference in adult gerbils and developing pups

We next asked whether there were any sex-based differences in the behaviors that we tracked (Fig 5). Across 3 cohorts we had 3 adult males, 3 adult females, 3 male pups and 9 female pups.

**Fig 5.**
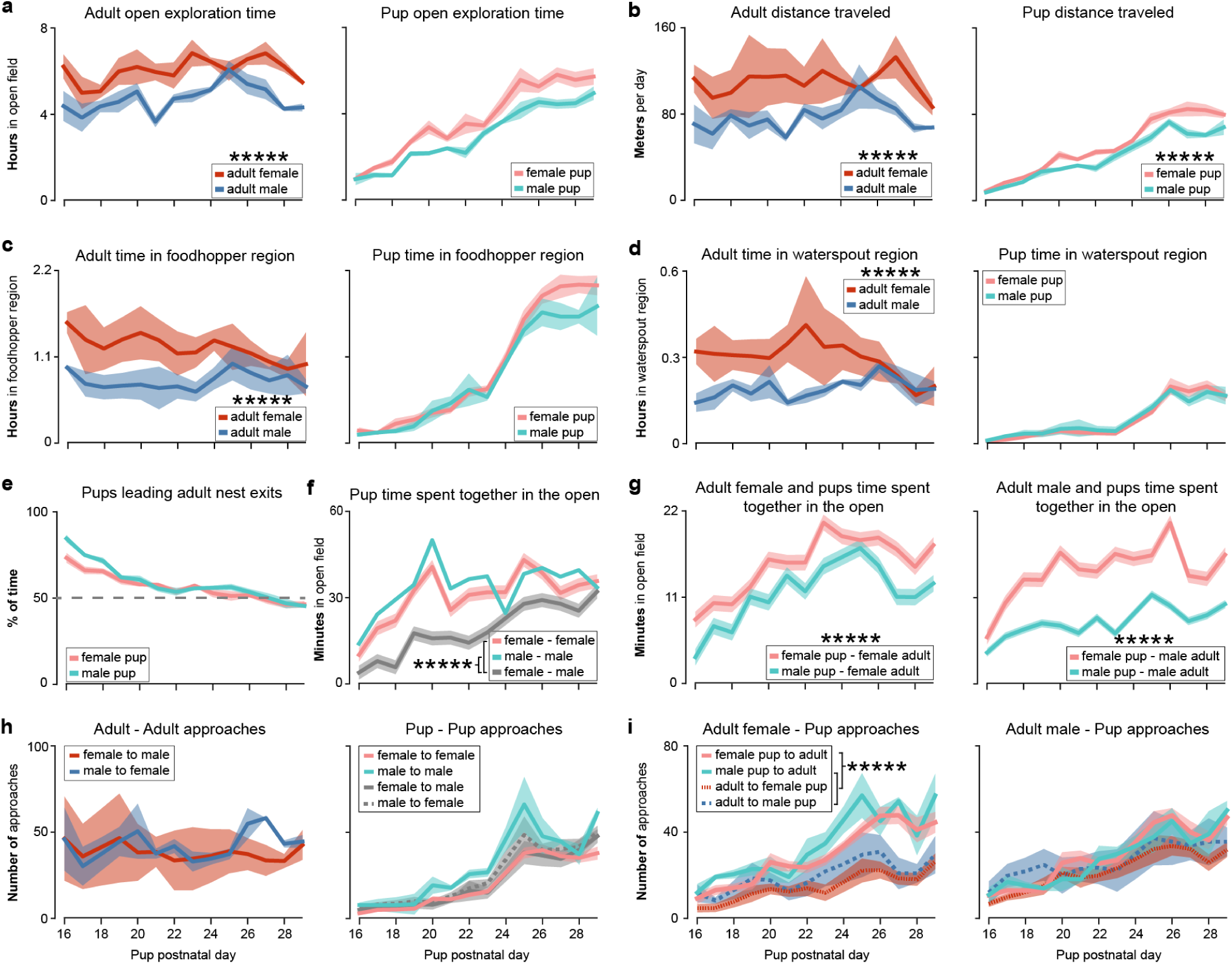
Sex differences in behavior. **(a)** The average (lines) and SEM (shading) of the time adult females (red) and adult males (blue) (left) and female pups (pink) and male pups (baby blue) explored outside of the nest. Asterisks indicate statistical significance on both t-test and 2 K-S test (e.g. p-value < 1e-05 = *****; see also Results and Methods). (**b)** Distance traveled by adults and pups. (**c)** Time spent inside the food hopper region. (**d)** Time spent inside the waterspout region. (**e)** The ratio of the number of times pups were followed by adults after leaving the nest. (**f)** Time spent in close proximity to each other in the open (pink: female with female; baby blue: male with male; gray: female with male). (**g)** Time the adult female (left) and adult male (right) spend in close proximity to female (pink) and male (baby blue) pups in the open. (**h)** The number of adult to adult approaches and pup to pup approaches. (**i)** Approaches between the adult female (left) and adult male (right) and pups.

*Female pups and female adults explore for longer periods and travel longer distances than males.* We quantified the amount of time spent exploring outside of the nest by sex for both adults and pups (Fig 5a). We found adult females on average spent 6.03 ± 0.90 hrs/day outside of the nest compared to males who spent 4.69 ± 0.83 hrs/day. For the 14 day recording duration, this yielded a total exploration time of 78.67 ± 5.05 hrs/cohort and 61.29 ± 1.44 hrs/cohort, respectively. The average daily difference in exploration between adult females and males was 1.35 ± 0.90 (t-test p-value: 5.51e-12; 2-sample K-S test against a shuffled distribution p-value: 1.67e-07; see also Methods). Similarly, female pups on average spent 3.73 ± 1.85 hrs/day outside of the nest compared to male pups who spent 2.94 ± 1.39 hrs/day outside of the nest, yielding a total exploration time of 48.96 ± 7.27 hrs/cohort and 38.25 ± 1.92 hrs/cohort, respectively. In contrast, female-pup to male-pup exploration time differences were smaller:-0.02 ± 0.87 (t-test p-value: 8.42e-01; 2-sample K-S test p-value: 9.54e-01; note; for pup exploration time the differences in Fig 5a were driven largely by our third cohort which did not have any male pups and was excluded from computation; see also Methods).

Similarly, adult females traveled on average 109.64 ± 31.14 meters/day compared to 76.44 ± 19.25 meters/day for adult males, resulting in total distance traveled of 1435.60 ± 294.64 meters for females and 1001.35 ± 82.39 meters for males for the duration of each cohort. The daily average distance traveled by females more than males was 33.20 ± 33.67 (t-test p-value: 1.54e-07; 2-sample K-S test p-value: 8.28e-05). Female pups also traveled on average longer 50.08 ± 27.66 meters/day than male pups 40.08 ± 21.32 meters/day, resulting in total distances traveled of 657.23 ± 79.69 meters and 523.77 ± 43.19 meters, respectively. The daily average difference in distance traveled by the female pups more than male pups was 7.41 ± 9.52 (t-test: 1.22e-08; 2-sample K-S test: 1.63e-07).

Overall, adult females spent on average 29% longer outside of the nest and traveled 43% more than males during the 14 day period we recorded. Female pups also traveled 25% more on average than male pups. These findings suggest a difference in adult male and female roles during periods of pup rearing.

*Female adults but not female pups spend more time in food hopper and water regions than males.* Adult females spent almost 100% more time than males in the food hopper area: average 1.40 ± 0.74 hrs/day in the food hopper area vs. males 0.73 ± 0.27 hrs/day in the food hopper region (totalling 18.42 ± 4.11 hrs/cohort and 9.55 ± 1.16 hrs/cohort for the duration of our recordings; Fig 5c, left). On average, females spent 0.43 ± 0.40 hrs/day more than adult males in the food hopper region (t-test: 2.76e-08; 2 sample K-S test: 2.96e-04). Female pups spent on average 0.83 ± 0.65 hrs/day and male pups 0.76 ± 0.62 hrs/day in the food hopper region, resulting in totals of 10.72 ± 1.31 hrs/cohort and 9.82 ± 0.36 hrs/cohort respectively (Fig 5c, right). On average, female pups spent 0.08 ± 0.23 hrs/day more than male pups in the food hopper region (t-test: 7.64e-03; 2 sample K-S test: 2.46e-01).

We found similar trends with respect to water spout region exploration with the adult female spending almost 80% more time in the water region than the adult male: average 0.27 ± 0.13 hrs/per day vs. males 0.15 ± 0.05 hrs/day (total times of 3.54 ± 1.36 hrs/cohort and 1.99 ± 0.36 hrs/cohort, respectively, over the entire recordings). On average, female adults spent 0.11 ± 0.13 hrs/day more than males in the water spout region (t-test: 6.36e-06; 2 sample K-S test: 1.50e-05). In contrast, female pups and male pups spent similar amount of time in the water spout region: female pups 0.06 ± 0.06 hrs/day, vs. male pups 0.07 ± 0.06 hrs/day (totals of 0.76 ± 0.21 hrs/cohort and 0.88 ± 0.25 hrs/cohort, respectively). (There were no significant differences between pup water spout behaviors).

We note that for the female adult-but not the male adult-the time spent in the food hopper region decreased 26% from P24 (1.31 ± 0.19 hrs/day) to P29 (1.00 ± 0.58 hrs/day) and the time spent in the water region decreased 41% from P24 (0.34 ± 0.12 hrs/day) to P29 (0.20 ± 0.10 hrs/day). This is consistent with a decreased role of the adult female in nursing the pups which underwent rapid development periods for food hopper proximity (beginning at P22) and water spout region (beginning at P24) (see Fig 2).

*Pup-pup pairwise foraging time shows intra-sex bias.* For social exploration behavior in pups, we found that female pups spent on average 30.48 ± 12.17 mins/day with another female pup and only 18.73 ± 10.91 minutes with male pups, i.e. 63% difference (Fig 5f). On average, female pups spent 5.53 ± 5.30 mins/day with other female pups more than with male pups (t-test: 2.28e-10; 2 sample K-S test: 4.89e-09). (We note that these values take into consideration the fewer numbers of male pups for this comparison; see Methods). While only a single cohort had two male pups, in that cohort the male pups also appeared to prefer within-sex time spent together (Fig 5f). On average, the male pups spent 18.66 ± 7.16 mins/day with each other more than with the female pups (t-test: 3.71e-07; 2 sample K-S test: 1.64e-03). Overall, this suggests the possibility of sex-based differences in social behaviors are entrenched from very early development-however, due to only a single cohort having multiple male pups, further experiments may be required to verify this.

*Adult to female pup social proximity time is significantly higher than proximity to male pups.* We found that adult females spent an average of 15.23 ± 6.09 mins/day with female pups outside of the nest and 11.59 ± 4.29 mins/day with male pups, a difference of 31% (Fig 5g). On average, the female adult spent 3.16 ± 4.91 mins/day more with female pups than male pups (t-test: 1.06e-06; 2 sample K-S test: 4.37e-04). The adult male had even more pronounced bias, spending 14.39 ± 6.00 mins/day with female pups and only 7.89 ± 2.97 with male pups, an 82% difference. On average, the adult male spent 5.57 ± 5.52 mins/day more with female pups than male pups (t-test: 3.57e-12; 2 sample K-S test: 1.07e-08). These results show the presence of a significant bias in socialization time with female pups over male pups, though it remains unclear what is driving such differences (e.g. male pups approach parents as frequently as female pups, but spend less time in the interactions; see also Fig 5i and Discussion).

*Sex-specific approach behaviors between pups and between adults and pups.* For adults, we found that males and females tended to approach each other a similar number of times per day, males approaching females: 42.17 ± 14.62 approaches/day and females approaching males: 38.10 ± 23.60 approaches/day (no statistically significant differences; t-test: 1.45e-01; 2 sample K-S test: 1.12e-01) (Fig 5h-left panel). In contrast, we found that male pup to male pup approaches occurred an average of 30.07 ± 20.64 times/day (cohort 2 data only), and female pups approaching female pups occurred 20.01 ± 16.66 times/day (Fig 5h, right). Our interpretation is that male pups tended to have shorter episodes of social interaction than female pups, suggesting a partially different type of social dynamic was at play (see Discussion).

We also found that adult females were approached significantly more by pups than the adult female approached the pups: average number of female pup to adult female approaches 29.07 ± 16.25 times/day and female adult approaches to female pups 14.46 ± 11.66 times/day, a difference of more than 100% (t-test: 3.55e-08; 2 sample K-S test: 2.46e-05). Male pups also tended to approach the adult female more: 34.05 ± 17.41 times/day while adult females approached male pups 19.57 ± 11.54 times/day, a difference of 74% (t-test: 5.67e-07; 2 sample K-S test: 2.59e-04). In contrast, the adult male approached female pups 21.96 ± 14.16 times/day, while female pups approached the adult male 28.86 ± 16.33 times per day, a difference of only ∼30 % (t-test: 1.01e-01; 2 sample K-S test: 1.83e-02) (Fig 5i-right panel). Similarly, the adult male approached male pups 26.52 ± 14.24 times/day, while male pups approached the male adult 27.76 ± 14.15 times/day, a difference of only ∼5% (t-test: 5.50e-01; 2 sample K-S test: 9.87e-01). These findings support that pups seek (i.e. approach) the adult female significantly more than the adult male but that this seeking behavior between pups and the adults is not symmetric in both cases.

In sum, we identify sex-dependent behaviors within both adults and pups and between the age classes. We find adult females spend more time in the open than males, travel longer distances, and spend more time in the food hopper and water spout regions. These findings may suggest a division of labor during critical development periods for pups-though it is unclear if such behaviors are present once pups have left the family. Within pups, we find that pups prefer socializing with their own sex substantially more than with the other sex-suggesting very early sex-based socialization preferences. Across the age classes, we find that both adult females and adult males tend to prefer socializing with female pups.

### The development of circadian rhythm based behaviors

We next evaluated the correlation between the animal behaviors and the day:night cycles. In particular, we sought to quantify significant changes at the onset of light cycles (Fig 6a).

**Fig 6.**
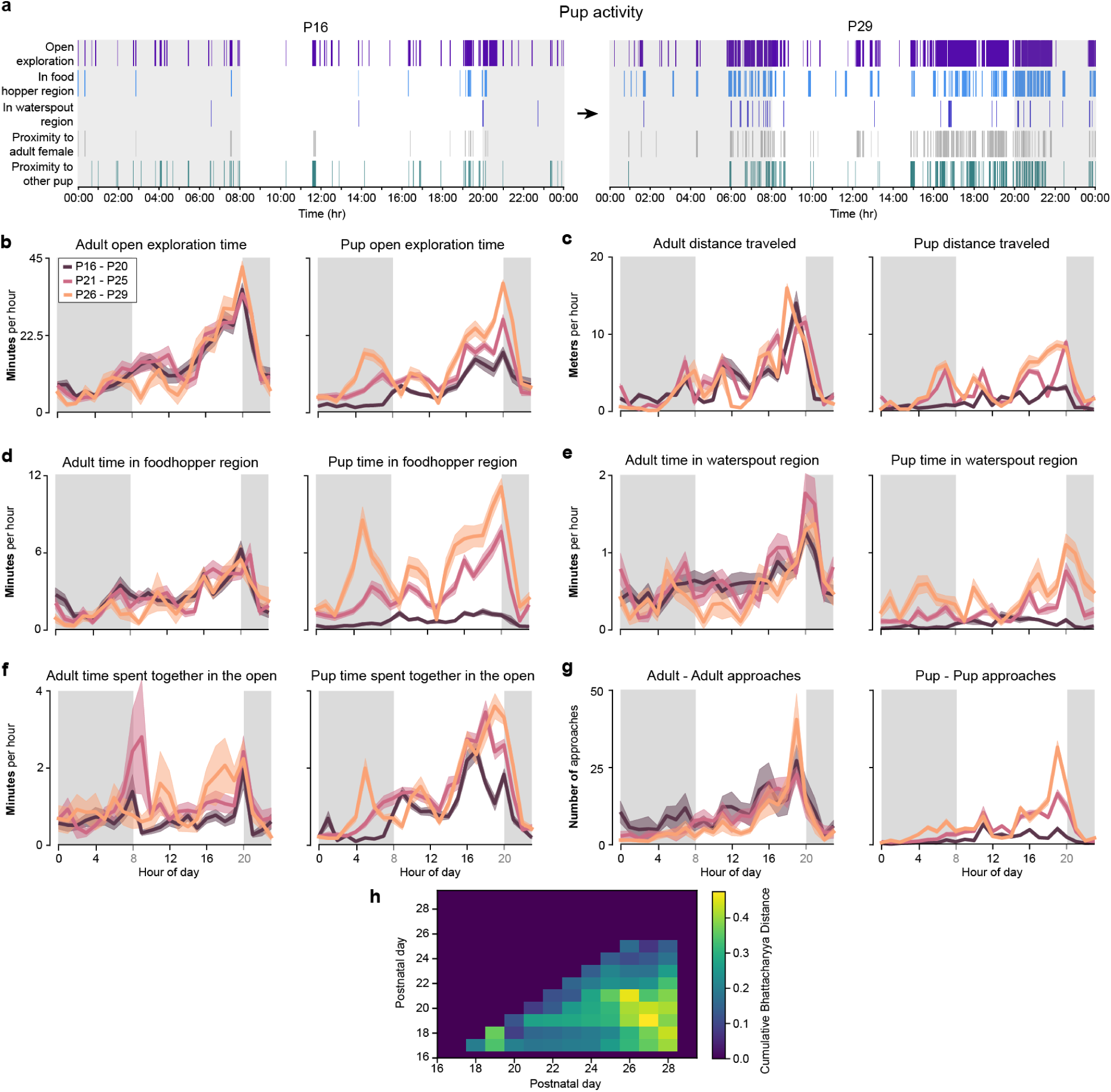
Circadian rhythms in adult and developing gerbils. **(a)** Behavior ethograms for one cohort showing substantial increases in the volume of behaviors between P16 and P29. **(b)** Average adult (left) and pup (right) open exploration time per hour (lines) and SEM (shading), across three developmental periods: P16 to P20 (dark purple), P21 to P25 (pink) and P26 to P29 (light orange). (**c)** Average adult (left) and pup (right) distance traveled per hour and SEM, across the three developmental periods. (**d)** Average adult (left) and pup (right) time spent in the food hopper region per hour and SEM, across the three developmental periods. (**e)** Average adult (left) and pup (right) time spent in the waterspout region per hour and SEM, across the three developmental periods. (**f)** Average time in proximity for adults with adults (left) and pups with pups (right) per hour and SEM, across the three developmental periods. (**g)** Average adult to adult (left) and pup to pup (right) approaches per hour and SEM, across the three developmental periods. (**h)** Cumulative Bhattacharyya distance (colors) for computing the largest circadian separation based on pup time in the food hopper region (see also Results).

*Diurnal and nocturnal behaviors.* Across most behaviors tracked we found that both adult and developing pups had similar day:night cycle exploration times or had slight preferences for day-cycle over dark-cycles (Fig 6). For example, the averages for adult foraging near the food hopper were 56.59% ± 0.53% during day cycle vs. 43.41 ± 0.46% for dark cycle while pup foraging was 59.6 ± 2.62% for day cycle and 40.4 ± 2.53% for night cycle (Fig 6d). In contrast, waterspout region averages were: adults 49.58 ± 0.7 % for day cycle and 50.42 ± 0.62 % for night cycle and pups: 48.8 ± 2.29% for day cycle and 51.2 ± 3.57 % for night cycle (Fig 6e).

*Genetically vs. environmental driven diurnal behavior structure.* Our observations of rapid development periods suggested approximately three different developmental stages: an initial period up to P19, a longer period up to P26 and a final stage from P26 to P29 during which all pup behaviors stabilized. We accordingly used this observation to initialize our analysis of development of circadian rhythms (though we also sought to optimize these transition periods as well; see Fig 6h below and Methods).

In adults, we observed both nocturnal and diurnal behaviors, with a slight bias towards diurnal behavior and a systematic increase in the amount of behavior during daytime and peaking between 19:00 and 20:00 (prior to the night cycle starting at 20:00). This ramping was present for most developmental periods in adults and for most behaviors studied including exploration time (Fig 6b), distance traveled (Fig 6c, food hopper proximity (Fig 6d), waterspout proximity (Fig 6e) and adult-adult approaches (Fig 6g, left panel). It was absent, or very weak in adult-adult proximity time (Fig 6f, left panel).

Similarly for pups, we also observed diurnal ramping in activity over the day cycle across most behaviors. However, this ramping up was present in all developmental periods only for exploration time (Fig 6b, right panel) and pup-pup time spent together (Fig 6f, right panel). For other behaviors, this ramping effect developed gradually in several behaviors: food hopper region exporation (Fig 6d), waterspout region proximity (Fig 6e), distance traveled (Fig 6c), and pup-pup approaches (Fig 6g, right panel). This suggests that the former behaviors (i.e. open exploration time and pup-pup social proximity) are likely driven by genetically encoded processes whereas the latter (food and water region exploration, distance traveled and pup-pup approaches) were driven by systems that develop later to detect valuable environmental and social interactions.

*The development of circadian behaviors.* We observed *circadian* behavior in both pups and adults for some, but not all, behaviors. In adults, *circadian* behavior was present in distance traveled (Fig 6c), weaker in water spout region proximity (Fig 6e), and mostly absent or very weak in food hopper region proximity (Fig 6d), social foraging time (Fig 6f) and adult-adult approaches (Fig 6g, left panel). (We note that our observation of a weak *circadian pattern* in waterspout region proximity is similar to Pietrewicz et al 1982). In pups, there were strong *circadian* behaviors in exploration time (Fig 6b), distance traveled (Fig 6c), foodhopper proximity (Fig 6d), pup-pup proximity time (Fig 6f, right panel). Waterspout region proximity had only a weak *circadian pattern* (Fig 6e) and there were few and small pre-dawn peaks in pup-pup approaches (Fig 6g, right panel). Importantly, *circadian patterns* developed over time for food hopper, pup-pup time spent together, exploration time and distance traveled-being mostly absent from early development (i.e. P16-P20). Interestingly, we note that qualitatively, pups appear to develop much more pronounced pre night-to-day cycle peaks generally around 4:00 and 5:00 than adults (night-to-day cycle change was at 8:00).

*Detecting the development of circadian rhythms.* We sought to computationally determine the transition from non-circadian to circadian rhythms-driven behaviors by agnostically determining the times of three developmental periods. In particular, we sought to determine the times of transition between: an initial period D1, a middle period D2 and a final period D3 (initialized as P16-P20, P21-P25 and P26-P29 above). We thus sought to determine the optimal two transition points, t1 and t2, determining the transition from D1 to D2 and D2 to D3, respectively, by maximizing the difference between the individual animal circadian trajectories in each of these periods (see also Methods). We used foodhoper region proximity as it was the most striking developmentally and we found t1=19 and t2=27, yielding three developmental periods of D1: P16-P19, D2: P20-P27, and D3: P28-P29. (Fig 6h). We note that as this method is based on initialization heuristics (e.g. assumes 3 developmental periods) different priors or assumptions may provide slightly different developmental windows.

## DISCUSSION

Using continuous recordings in an enlarged home-cage paradigm, we sought to quantify the development of autonomous and social behaviors in gerbil pups-described together as the development of agency-across a highly dynamic developmental window, i.e. P15-P30, during which gerbil pups go from being almost exclusively in the nest to being nearly independent with respect to many developmental measures. We employed machine vision tools and developed methods for quantifying a set of unique developmental trajectories. Rodent families have been studied both in natural environments to establish ground truth measurements (Ågren 1984; Ostermeyer & Elwood 1984; Hurtado-Parrado et al 2015), and in simplified laboratory enclosures to conduct mechanistic experiments (Lahvis 2017; Bordes et al 2023). Laboratory studies generally exploit brief, stereotyped behaviors in which one or two subjects are tested in a foreign environment. While this standardizes and simplifies the experimental design, it can disrupt diurnal cycles (e.g., feeding, sleep), perturb arousal or motivational states (e.g., human sensory stimuli; Sorge et al 2014), and introduce unanticipated variation (Crabbe et al 1999; Chesler et al 2002; Richter et al 2009). Some quantitative descriptions of rat family behavior in home cage environments have been attempted, but relied on human scoring of videos (e.g. Thiels et al 1990).

*Pup behaviors develop along distinct time courses.* For solitary behaviors, such as time spent exploring and distance traveled per day, we observed a gradual increase between P16 and P26, when pups reached adult-like levels. Food proximity behavior displayed two phases, increasing gradually at first, but escalating dramatically from P22-P26, and stabilized at 200% of the adult average. In contrast, water proximity behavior emerged slowly, increasing substantially only on P24 and reaching ∼80% the adult average by P26. Taken together, these results suggest the prolonged development of exploration behaviors (e.g. distance traveled and exploration time), but a distinct value-driven emergence of food and water exploratory behavior. The timing of the rapid development periods for food exploration (P22) and water exploration (P24 onwards) coincide with the adult female decreasing time spent in the food hopper region (26% decrease from P24) and the time spent in the water region (41% decrease from P24). These findings provide independent confirmation that the pups’ increase of food hopper and water spout proximity were driven, in part, by consumption behaviors and a decreased dependence on the adult female.

With respect to social behaviors, we found distinct trajectories for passive (e.g. remain in nest), interactive (e.g. social proximity) and active (e.g. approach) behaviors. Pups stayed in the nest approximately 20% more than adults at P16, and only reached adult levels at ∼P24. However, when pups did leave the nest, adults tended to follow them during this period (Fig 3b). These behaviors confirm that pup exploration of the environment increased gradually, likely coinciding with maturation of critical sensory-motor systems over this period (Tan et al 2017). With respect to social proximity time, we found that P16 pups tended to spend an equivalent amount of time with each other or adults (<20% of the adult-adult average). However, this pattern diverged significantly by P19 with pup-pup proximity time rising rapidly and stabilizing above the adult average (Fig 3d). In contrast, pup-adult proximity stabilized at ∼60% of the adult average. Our interpretation is that pups value social interactions with other pups from an early age. Lastly, P16 pups tended to approach each other < 10% of the adult average compared to approaching adults ∼25% of the adult average. This difference persisted until P24-25 when both approach behaviors stabilized to adult-like levels. To reconcile this with the larger time that pups spend together, we suggest that pups approached adults more often during early development, but remained within proximity for shorter periods of time.

*Behavior development is plausibly supported by the maturation of spatial-cognition and motor systems.* A systematic finding in our study is that all behaviors commence at near zero levels relative to adults at P16, and mature with distinct time courses between P16 to P26. This period coincides with the maturation of neural pathways that subserve spatial memory especially in the hippocampal and medial-entorhinal-cortex axis (HPC-MEC). In mice, there are several systems that develop during this period: formation and maturation of head direction cells (P10-P20), place cells (P14-P40), boundary responsive cells (P16-17) and grid cells (P20-P21) (Tan et al 2017). In rats, hippocampus dependent learning also has been shown to commence around P18 for water maze learning (e.g. visual platform-P17-20-and hidden platform-P19-21-learning), and T-maze learning (P19-P42) (Wills et al 2013). In fact, social encoding of family membership may also reside in the hippocampus (Boyle et al., 2024). The behaviors that we observed are not closely correlated with the onset of sensory transduction. The onset of transduction displays a similar sequence in most vertebrates, progressing from somatosensory to vestibular, olfactory, auditory and, finally, to the visual system (Gottlieb 1971). In mice, these systems emerge between birth and P12. However, a prolonged development of sensory processing has been demonstrated for many mammals (Sanes et al 2019). Therefore, one plausible interpretation is that the highly dynamic and specific periods of development that progress from a minimal state to near-adult levels between P16 to P29 are driven, in part, by maturation of HPC-MEC axis.

A second plausible interpretation is that the changes and stabilization we observe arise as the motor cortex matures. In particular, the motor cortex is still relatively immature (e.g., as compared to sensory cortices), with an estimate that M1 starts producing a significant outflow only after postnatal day 20 (Blumberg and Adolph 2023). Cortical motor outflow appears to begin at ∼P24 in rats (Young et al 2012; Ramanathan et al 2015; Singleton et al 2021), and there is a significant increase in stimulation-related motor activity after this age. For example, M1 activity recorded in vivo in P19-23 rats has yet to exhibit activity before wake or sleep movements (Dooley et al 2021). Therefore, an increase in exploration may also be tied to the maturation of the motor cortical pathway.

*Possibility of discrete stages of behavior development.* Our findings suggest three developmental periods: the first is approximately three days (P16 to P19), the second is more prolonged (P17 to P26), and the third begins later (P20-26). While it is intriguing to try and match these identified behaviors with specific developmental programs in neuronal maturation, our evidence only supports a very broad correlation with early developing behaviors likely driven by the early capacities of sensory systems, middle and prolonged periods of development driven by motor outflow and spatial systems (e.g., the HPC-MEC axis), and later behaviors being driven by increasingly sophisticated social-based factors that likely require additional CNS maturation. Importantly, we note that these types of sequential developmental behavior stages are also present in the sequential maturation of entorhinal-hippocampal networks which go from nearly 0% mature at P8-P14 to > 80% mature by P26 (e.g. Donato et al 2013 on mouse maturation).

*Sex-differences in behavior in environment explorations.* In adults, we found that females explored for longer periods (29%) and traveled further (43%) than males, consistent with other studies (Roper 1976; Ru-yung & Shao-liang 1984). Similarly, females spent almost double the amount of time as males in the food hopper region and 80% more in the water region than males. Furthermore, there was a significant decrease in these differences beginning ∼P25, consistent with pups relying on mother care and nursing around this time. In pups, we did not find a significant effect of sex on exploration time, but there was a small significant difference for distance traveled. Taken together, these findings support other behavioral studies of a possible sex-based bias in animal foraging times possibly present very early in development prior to sexual maturation.

*Sex-differences in social interactions between pups.* We found several significant differences in pup-adult and pup-pup interactions based on pup sex. In social exploration, we found that female pups spent 30% more time with other female pups than with male pups. Similarly, we found male pups spent almost 100% more time with each other than with female pups (note: this came from a single cohort that had 2 male pups). Similar to above, we point out that these sex-based differences in pup-pup interactions appear to be present from very early development.

*Sex-differences in social interactions between pups and adults.* Adult females spent more time with female pups (31%) than with male pups, and the adult males spent 82% more time with female pups than with male pups. In contrast, we found that both female pups and male pups tended to approach the female adult significantly more than she approached them (> 100% and 74% respectively). This asymmetry was not present in adult male to pup approaches. These findings suggest a process by which female pups may be more interested in socializing with adults than male pups, with both pups being significantly more interested in proximity to the female adult than the male adult. While this finding is supported by the need for pups to nurse in early development, we did not observe a decrease in this asymmetry over development, suggesting a more persistent bias being present at least until the end of our analysis period (P29).

*Circadian rhythms in adults.* Our paradigm enabled us to track the circadian rhythm effects on adult behaviors and the development of circadian rhythm correlated behaviors in pups. Studies on behavior and circadian rhythms in gerbils indicate that light conditions can affect behavior (Pietrewicz et al 1982; Probst et al 1987; Refinetti & Kenagy 2018), with a crepuscular pattern for most behaviors except drinking (Pietrewicz et al 1982). In our study, we found adults were active during both day and dark cycles, with more behaviors during day time and a substantial ramping of activity prior to night cycle onset. This ramping up is reminiscent of crepuscular behaviors in field studies, but not completely identical. We also observed diurnal behavior patterns only in the amount of distance traveled with other behaviors showing weak or no crepuscularity. We note that increased activity during the day is consistent with findings of greater maternal attention to pups during the day than at night (Grota & Ader 1969; Levin & Stern 1975).

*Development of circadianity in pups.* Beginning at ∼P26, pups began to display diurnal patterns in all behaviors with peaks at night cycle onset. However, in contrast to adults, these older pups also displayed substantial diurnal ramping behaviors as light-change switches approached across most behaviors (note: pup behavior was weak in the pup-pup approach behavior-similar to adults). This suggests that pups, unlike adults, had stronger drives to increase their activity pre day cycle changes than adults. Considering all periods of development, however, we found that some behaviors such as open exploration time and pup-pup time spent together changed much less between early and late development compared to food hopper region and waterspout proximity. We sought to use the heterogeneity in food hopper proximity in particular-as it showed the largest differences over time-to agnostically detect different developmental stages for circadian rhythms, and identified P16-19, P20-27 and P28-29 as the most likely partitioning of development into three stages.

*Limitations of our work and directions for future studies.* Our enriched and enlarged home cage environment enabled the tracking of several types of behaviors. However, our environment did not capture other behaviors observed in the wild such as gerbils’ increased tendency for exploration towards novelty (Osborne 1977) and towards “open-arms’’ of mazes (compared to rats and mice; Rico et al 2016). Gerbils also show increased unpredictability in their escape behaviors than other rodents (Moore et al 2017), and it would be interesting to observe this escape behavior dynamics by simulating predators within our environment. Beyond home cage environments, we propose that contemporary machine learning methods are improving substantially and it may be possible to carry out similar studies in “limited size” environments such as scientifically-purposed barns (Lahvis 2017).

*Understanding development from the micro to the macroscale.* Our study supports the viability of studying rodent behavior across orders of magnitudes of temporal resolution during highly dynamic periods of behavior and development in family groups of rodents. Our findings reveal novel time courses for development in rodents, including dynamic periods of behavior acquisition, sex-based behavior differences, and the development of circadianity. We view that increasingly ethological (e.g. group and family housed animals) and naturalistic (e.g. larger home enclosures with increased enrichment opportunities) behavior paradigms in the laboratory environment is an adequate and complementary method for the characterization of behavior and CNS development in health and disease.

## METHODS

### Experimental animals

Three gerbil families (Meriones unguiculatus, n=6 per family: 2 adults, 4 pups) were used in this study (Charles River). All procedures related to the maintenance and use of animals were approved by the University Animal Welfare Committee at New York University, and all experiments were performed in accordance with the relevant guidelines and regulations.

### Audio recording

Four ultrasonic microphones (Avisoft CM16/CMPA48AAF-5V) were synchronously recorded using a National Instruments multifunction data acquisition device (PCI-6143) via BNC connection with a National Instruments terminal block (BNC-2110). The recording was controlled with custom python scripts using the NI-DAQmx library (https://github.com/ni/nidaqmx-python) which wrote samples to disk at a 125 kHz sampling rate. In total, 13.084 TB of raw audio data were acquired across the three families. For further analyses, the four-channel microphone signals were averaged to create a single-channel high-fidelity audio signal.

### Home cage enclosure

We used an enlarged home enclosure (WxLxH: 35.6cm x 57.2cm x 38.1cm) with *ad lib* access to a food hopper and a water source.

### Video recording

We recorded video using an overhead camera, FLIR USB blackfly S (FLIR USA), at 24 frames per second. We recorded individual periods of approximately 20 minutes (cohort 1) or 60 minutes (cohorts 2 and 3) following which video and metadata were saved for offline use. The saving period resulted in approximately 10% downtime between recordings. Continuous video recordings were obtained from an overhead camera (24 fps), and audio recordings were obtained from four ultrasonic microphones (125 kHz sampling rate).

### Night:day cycle

Daytime began at 08:00 and night began at 20:00 during which an infrared light was activated.

### Animal shaving patterns

We used a distinct shaving pattern across all our cohorts (as described in Fig 1b). We opted for use of shaving patterns instead of fur bleaching, implanted trackers or body piercing as we viewed shaving as a less intrusive method.

### Tracking-Features

We manually labeled 1000 frames using the SLEAP (Pereira et al., 2022) labeling workflow. The skeleton consisted of 6 nodes: nose, spine1, spine2, spine3, spine4, and spine5; and 5 edges: nose to spine1, spine1 to spine2, spine2 to spine3, spine3 to spine4, and spine4 to spine5. Additionally, we labeled each instance with the animal’s identity class as female, male, pup1, pup2, pup3, and pup4 to enable tracking unique identities. Due to idiosyncratic differences in shaving patterns, lens distortion and light conditions, each cohort and light condition (i.e. day and night) was labeled and trained individually, resulting in 6 feature models. The final results were trained using SLEAP bottom-up ID models that achieved 97% accuracy of adults and 91% for pups.

### Track clean up-features

We developed an ID-switch error detection method to detect and correct some of the SLEAP ID-switch errors. We defined ID switch errors as those for which track segments were swapped between two animals. These errors were identified by evaluating sequential segments of tracks produced by SLEAP and determining whether (i) a large jump in track location occurred; and (ii) whether another segment fit better (usually within < 5 pixels of distance). We additionally implemented an interpolation method that connected SLEAP segments < 3 cm apart that belonged to the same animal. A custom python pipeline was developed to carry out this correction.

### Tracking-Huddles

We manually labeled 1000 frames using the SLEAP (Pereira et al., 2022) labeling workflow. The skeleton consisted of 1 node: huddle; and no edges. We identified a huddle as a group of 2 or more animals grouped together in a stable location, which were usually places that animals huddled in for the entire recording period or several sequential days. Due to idiosyncratic differences in shaving patterns, lens distortion and light conditions, each cohort and light condition (i.e. day and night) was labeled and trained individually, resulting in 6 huddle models. We additionally implemented an interpolation method that connected SLEAP segments for huddles that were less than 30 seconds apart.

### Bedding changes and exclusion of time points

We provided an interpolated measure of behavior on days of bedding changes. Briefly, on cohorts 1 and 2 we averaged the behaviors between P23 and P25 instead of using P24. On cohort 3 on P18 we averaged the behaviors between P17 and P19 instead of using the P18 day as we observed a substantial difference in the behavior just for that day.

### Cross-validation of behavior tracking

We implemented 10-fold cross-validation to evaluate the precision of SLEAP identity tracking against a human annotator. Briefly, for all feature NNs we split our human labeled frames dataset (1000 labeled frames) into 950-training and tested on the hold out of 50 frames. We repeated this 10 times by randomizing the training set frames and computed the average identity accuracy errors across adults and pups.

### Convex hull overlap metric

We defined overlap volume between behaviors as the intersection between the convex hull generated by individuals in each type of behavior relative to the other behaviors. We computed the overlap as the intersectional volume of the first 3 principle components (using the python vedo package) or when this was not possible due to insufficient points to form a polyhedron-the first 2 principle components. We viewed the overlap volume as better at capturing and representing the inter-behavior similarity by providing a metric that quantifies when some of the individual behaviors were not distinguishable between behavior types. We note that comparing the individual behaviors at each time point was not practical due to only 3 samples in the behavior of the adult female and male and male pup. We also note that we did not use more than 3PCs (or the full dimensionality of the distributions) to compute these similarity metrics as algorithms to compute multidimensional overlapping volumes are not available and due to the curse of dimensionality we would likely get zero or vanishingly small overlaps.

### Rapid development and stabilization periods

We defined a rapid development period as Pdays that had distributions that were statistically significant than the previous 2 Pdays. We thus pooled behaviors in groups of 2 days and compared them following groups. We used two days as a more conservative way to account for higher variability-than applying smoothing or other filtering techniques on the raw data. We used a 2 sample Kolmogorov-Smirnov test as a more conservative measure of significance. For clarity, at each Pday, we pooled data from individuals (e.g. 12 pups or 6 adults) and computed the statistics across these values. We used the python package *statsmodels* method *sm.stats.multipletests* for the Benjamini-Hochberg multiple hypothesis testing adjustment and reported pvalues < 0.05 as statistically significant.

### Heuristics-based behavior classification

We implemented heuristics to compute the location of animals near the food hopper or waterspout and each other. For the food hopper we used an ROI of size 19.50 × 10.50 cm. For waterspouts we used an ROI of size 8.50 × 8.00 cm. For inter-animal proximity we used a distance of < 5 cm separation and a minimal proximity time > 200 ms. Animal locations were computed as the median centroid over (x,y) locations. For control and comparison to ROIs in food and water spout regions we also selected ROIs in random locations of the cage. We found that ROIs that were randomly placed had little occupancy (limiting statistical analysis) and did not have the dynamics of ROIs in the food and water spout regions.

### SimBA-based behavior classification

We defined an approach as an animal moving towards another-beginning when the initiator starts moving and ending when it stops moving (at most one gerbil length away from the other) or makes physical contact. We manually annotated 300 approaches (4578 frames) in 22 videos and implemented a modified version of SimBA (Nilsson et al 2020) for classifying pair-wise animal approach behaviors. We computed the threshold for approach behavior as the 99% percentile of the prediction distribution. We verified that detected approaches matched with qualitative approaches.

Descriptive statistics of animal movements, angles, and distances in sliding time-windows were computed using runtime optimized methods available through the SimBA API and used in a downstream random forest classifier. We randomly sampled annotated frames from separate events in training and test set to reduce time-series data leakage. Within the training set, we further balanced the data by under-sampling the number of non-event observations to a 1:1 ratio of the event observations. Classifiers were validated and optimal discrimination threshold determined by evaluating performances on novel held-out videos excluded from the training and test sets.

### Multi-animal social state networks

We developed an approach to quantify unique social configuration states and their dynamical transitions. We first discretized the cage space into 5 x 4 squares of approximately 11cm x 9cm each-yielding a 20 Dimensional state space. At each time point in our recordings we computed the occupancy of each location as the number of animals at that location. This yielded an N_videoframes_ x 20D matrix which in general had a few hundred or thousand unique 20D vectors per PDay. Each 20D vector thus contained integers from 0 (no animal in the location) up to 6 (i.e. all animals were at the grid location). Connectivity matrices for each PDay were then computed by connecting each of the videoframe nodes to the one occurring subsequently. We used the connectivity matrices to generate 14 networks for each PDay of our study and for a variety of animal configurations. For clarity, we computed 5 types of networks: single pups (i.e. 1 animal only), pairs of animals (the 2 adults and all pairwise combinations of pups), all-pups (i.e. all 4 pups) and all-animals (all 6 animals).

Fig 4a,b) (see also Methods). This enabled us to generate a network-based analysis where each time point in our recording generated a 20 Dimensional (20D) social state vector and we could generate a network connectivity matrix by connecting all social states that occurred sequentially (Fig 4c; see also Methods). The network connectivity matrix enabled us to generate graphs that represented the social state dynamics at each PDay in our cohorts (Fig 4d).

### Inter-sex statistical differences tests

We used a two sided student t-test using the SciPy python library to evaluate the significance of the difference between our distribution (usually the difference between two groups-see below) and a zero-mean distribution obtained by shuffling the data. Additionally, we used a 2 sample K-S test from the SciPy library to compare the original distribution with a shuffled version where the identity of the animals was randomized. For example, for computing the distribution of distance traveled by adult females vs. adult males, we subtracted the distance traveled by each male from distance traveled by each female on each day of each cohort. This provided a day-wise difference in the behavior of each animal. For the shuffled version, we did the same but randomly switched the sex identity of the animals, i.e. subtracting male from female and female from male at random. For pups we only used cohorts 1 and 2 as they were the only ones with male pups. In cohort 1, there were 2 pups and there we computed the average male behavior as the reference behavior and subtracted it from the individual female pup behaviors to get our distributions. We did not adjust for multiple hypothesis testing as only a single statistical test was performed within each behavior class. We also reported the pvalues in full and drew conclusions from those that were much lower than 0.05 (i.e. Bonferonni correction for 32 tests which are all tests carried out across all behaviors would lower the requirement for significance by ∼1.5 orders of magnitude; and Benjamini-Hochberg correction would lower this requirement by even less).

## Data availability

All datasets including raw behavioral videos, trained SLEAP neural networks and pose estimation tracks are available and will be deposited with Dryad. Video datasets are between 5-10TB in size per cohort and contain tens of thousands of files including original resolution videos, compressed videos, SLEAP output tracks, and SimBA training data and inference files.

## Computer Code

Software processing pipelines are publicly available and include SLEAP (https://github.com/talmolab/sleap), SimBA (https://github.com/sgoldenlab/simba) and custom postprocessing code available at https://github.com/catubc/gerbil.

## Acknowledgements

Supported in part by NIH R01DC020279 (DHS). We thank Garet Lahvis, Mark Blumberg, Kenny Kay and Paola Cerrito for their helpful suggestions or comments on the manuscript.

## Author contributions

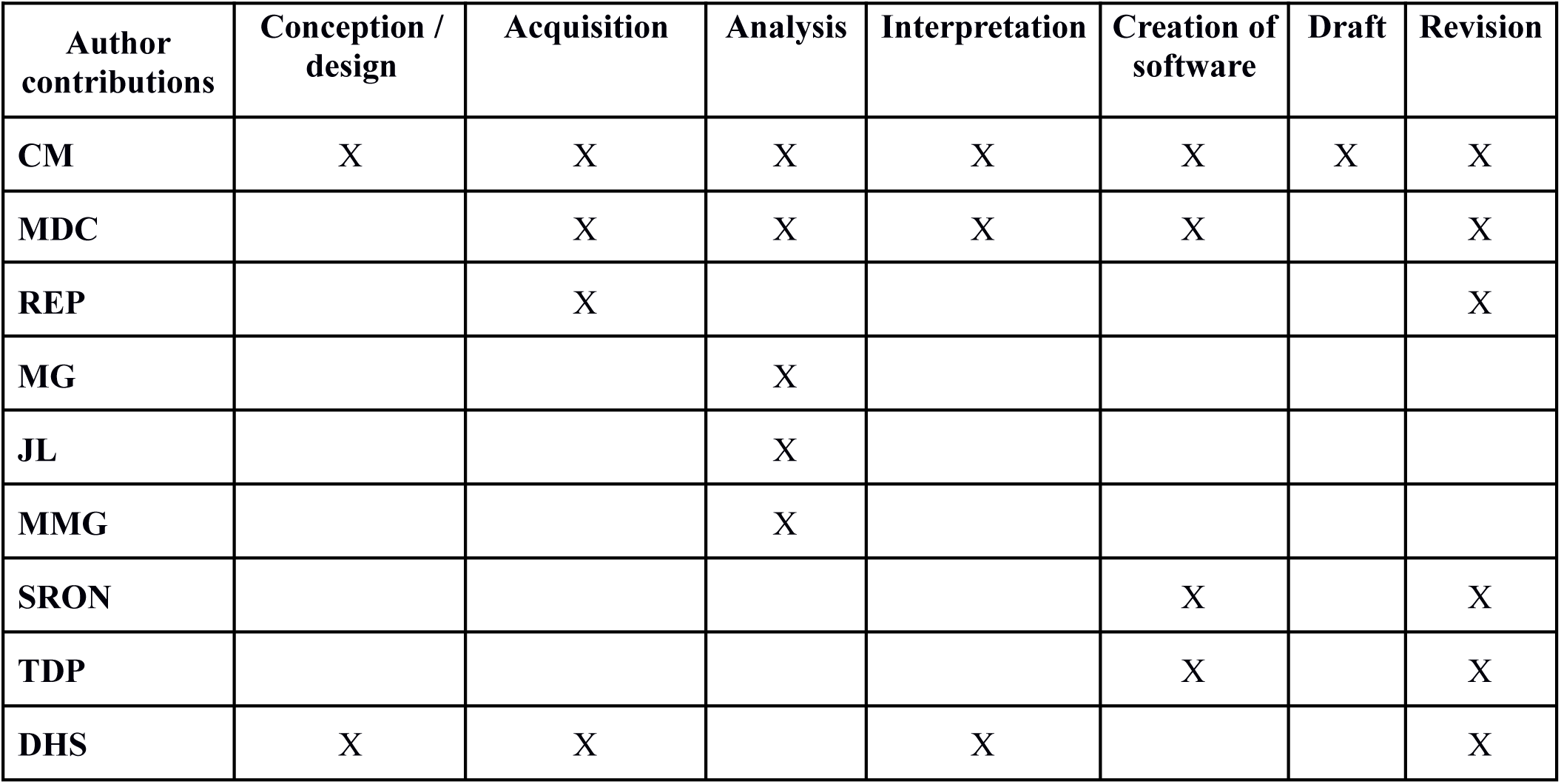

